# Migraine Attacks as a Result of Hypothalamic Loss of Control

**DOI:** 10.1101/2020.11.19.390104

**Authors:** Anne Stankewitz, Leonie Keidel, Mathias Rehm, Stephanie Irving, Stephan Kaczmarz, Christine Preibisch, Viktor Witkovsky, Claus Zimmer, Enrico Schulz, Thomas R Toelle

## Abstract

To assess the natural trajectory of brain activity over the migraine cycle, we assessed (1) the cerebral perfusion and (2) the hypothalamic connectivity during spontaneous headache attacks with follow-up recordings towards the next migraine attack. Using pseudo-continuous arterial spin labelling and resting-state functional magnetic resonance imaging, 12 migraine patients were examined in 82 sessions. We detected cyclic changes of brain perfusion in the limbic circuit (insula, hippocampus, and nucleus accumbens), with the highest perfusion during headache attacks. In addition, we found an increase of hypothalamic connectivity to the limbic system over the interictal interval, then collapsing during the headache. Our data provide strong evidence for the predominant role of the hypothalamus as a zeitgeber for generating migraine attacks. Our findings suggest that migraine attacks are the result of the hypothalamus losing control over the limbic system.

## Introduction

Migraine is a complex neurological disorder that is affecting approximately 12% of the population (*1*). Recurring headache episodes are often the most disabling symptom reported by migraineurs, however, migraine attacks are more than just a headache. Typical migraine attacks span 4 to 72 hours, containing up to four phases: premonitory phase, migraine aura, headache phase, and the postdrome (*2*). Although the pathophysiology is not yet fully understood, strong evidence has emerged that both the peripheral trigemino-vascular system (*3*) and the central nervous system contribute to the genesis of the disease (*4–6*).

In particular, the hypothalamus has been highlighted as a major player in initiating, maintaining, and remitting migraine attacks (*7, 8*) via its strong connections to various cortical and subcortical brain regions (*9, 10*). The hypothalamus is known to regulate endogenous circadian and circannual rhythms and to maintain internal homeostasis by controlling the endocrine and the autonomous nervous system (*11, 12*). The clinical features of migraine underline the hypothalamic contribution to the pathogenesis of the disease, i.e. the cyclic behaviour of attacks, prodromal symptoms (yawning, fatigue, and mood changes), autonomous symptoms (nausea and vomiting), as well as trigger factors, such as certain hormones and psycho-physiological stress (*7, 13*).

The first observations of an increased cerebral blood flow bilaterally in the hypothalamus during spontaneous headache attacks have been made with positron emission tomography (PET) (*14*). However, due to the unpredictable onset of migraine attacks, the investigation of the cyclic behaviour of hypothalamic activity is challenging and the group size of studies is therefore often limited (*15–17*). To cut short the natural but uncertain course of the migraine cycle, some studies applied nitroglycerin to patients in order to trigger headache attacks (*18, 19*). Implementing such a stimulation design, Maniyar et al. did not observe hypothalamic alterations at the time of the headache, but during the premonitory phase prior to the nitroglycerin-induced headache (*19*).

Likewise, functional magnetic resonance imaging (fMRI) studies have applied painful stimulation in order to stress the trigeminal system (*17*). Investigating the effects of external stimuli instead of the more relevant endogenous pain is a known methodological limitation of functional imaging, as the change of cortical activity can only be determined in contrast to a baseline or to a functional control condition. Recently, a longitudinal fMRI study observed increased hypothalamic activity in response to painful stimulation during the 48 hours prior to the beginning of headache attacks. By contrast, the hypothalamus was not active during the migraine attack (*17*). Using the same trigemino-nociceptive and olfactory stimulation design, previous studies did not find any hypothalamic alterations neither during (*15, 16*) nor within the last 72h prior to spontaneous headache attacks (*16*).

Other fMRI studies have focussed on hypothalamic functional connections, finding stronger hypothalamic connectivity in response to painful trigeminal stimulation in migraineurs during headache attacks compared to the pain-free interval (*20, 21*) and during the prodromal phase (24h prior to spontaneous migraine headache) (*20, 22*). However, the very same methodological limitations apply; the natural cycle of the migraine is not being measured.

The present study was designed to circumvent such limitations by exploring the natural trajectory of the endogenous pathological rhythmicity in migraine over an entire migraine cycle by combining two important aspects:

*First*, we used a longitudinal design to assess the pathological brain processes over the entire migraine cycle in a group of migraineurs at several time points: we started from the ictal phase of a migraine attack and recorded the disease-related cortical processes throughout the interictal phase towards the next migraine attack.

*Second*, we followed the trajectory of the endogenous pathological cortical processes, untriggered and unstimulated, over its natural cycle. The pseudocontinuous arterial spin labelling (pCASL) technique allows non-invasive measurement of cerebral blood flow (CBF), which is commonly used as a proxy for neuronal activity. Seed-based resting state functional MRI was specifically used to trace rhythmic changes in hypothalamic connectivity, which is considered to trigger the attacks.

## Results

### Cerebral blood perfusion (pCASL)

To determine the cyclic changes of the migraineurs’ brain activity between two migraine attacks, we computed a linear mixed effects (LME) model and analysed how the cortical perfusion (pCASL) is dependent upon the time point within the migraine cycle.

In the first part of our analyses we were testing for a “drop” of cortical activity during headache attacks (“reset mode”). We found that the superior parietal cortex showed significant cyclic activity, with lowest cortical perfusion during the headache that increased steadily afterwards (Table 1). There was no cyclic activity decreasing from the day of the attack for cortical perfusion.

**Table 1.**
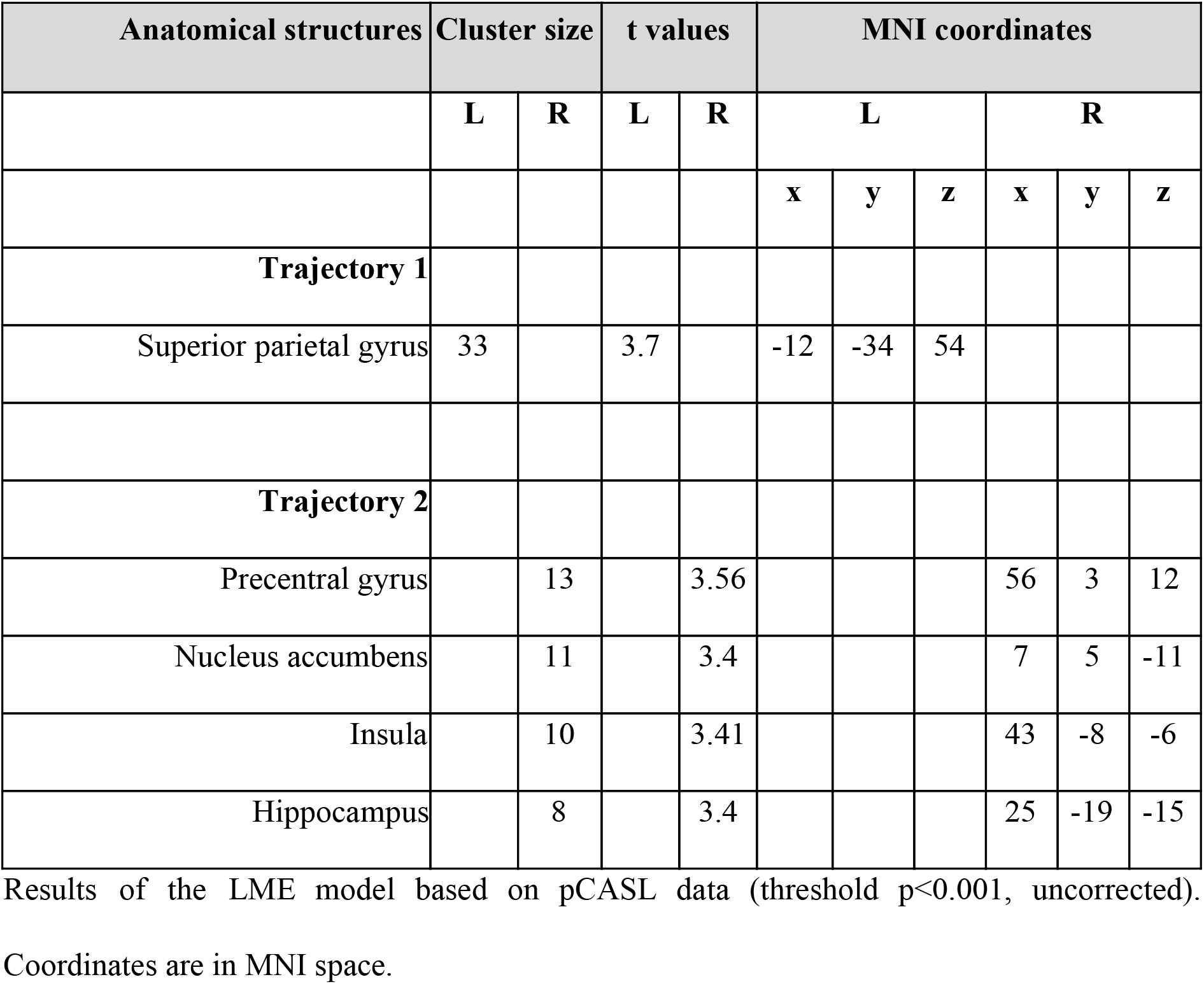
Brain perfusion data over the migraine cycle.

| Anatomical structures | Cluster size |  | t values |  | MNI coordinates |  |  |  |  |  |
| --- | --- | --- | --- | --- | --- | --- | --- | --- | --- | --- |
|  | L | R | L | R | L |  |  | R |  |  |
|  |  |  |  |  | x | y | z | x | y | z |
| **Trajectory 1** |  |  |  |  |  |  |  |  |  |  |
| Superior parietal gyrus | 33 |  | 3.7 |  | -12 | -34 | 54 |  |  |  |
| **Trajectory 2** |  |  |  |  |  |  |  |  |  |  |
| Precentral gyrus |  | 13 |  | 3.56 |  |  |  | 56 | 3 | 12 |
| Nucleus accumbens |  | 11 |  | 3.4 |  |  |  | 7 | 5 | -11 |
| Insula |  | 10 |  | 3.41 |  |  |  | 43 | -8 | -6 |
| Hippocampus |  | 8 |  | 3.4 |  |  |  | 25 | -19 | -15 |

Results of the LME model based on pCASL data (threshold $p < 0.001$ , uncorrected).
Coordinates are in MNI space.

Next, we tested for cortical activity that is changing towards the migraine attack and reaching its climax during the attack (“pain mode”). We observed increasing blood perfusion with a maximum during the attack in the right insular cortex, the right hippocampus, the right nucleus accumbens, and the right precentral gyrus (Table 2 and Fig. 1). We did not find any cortical area that showed declining perfusion during the migraine attack.

**Fig. 1.**
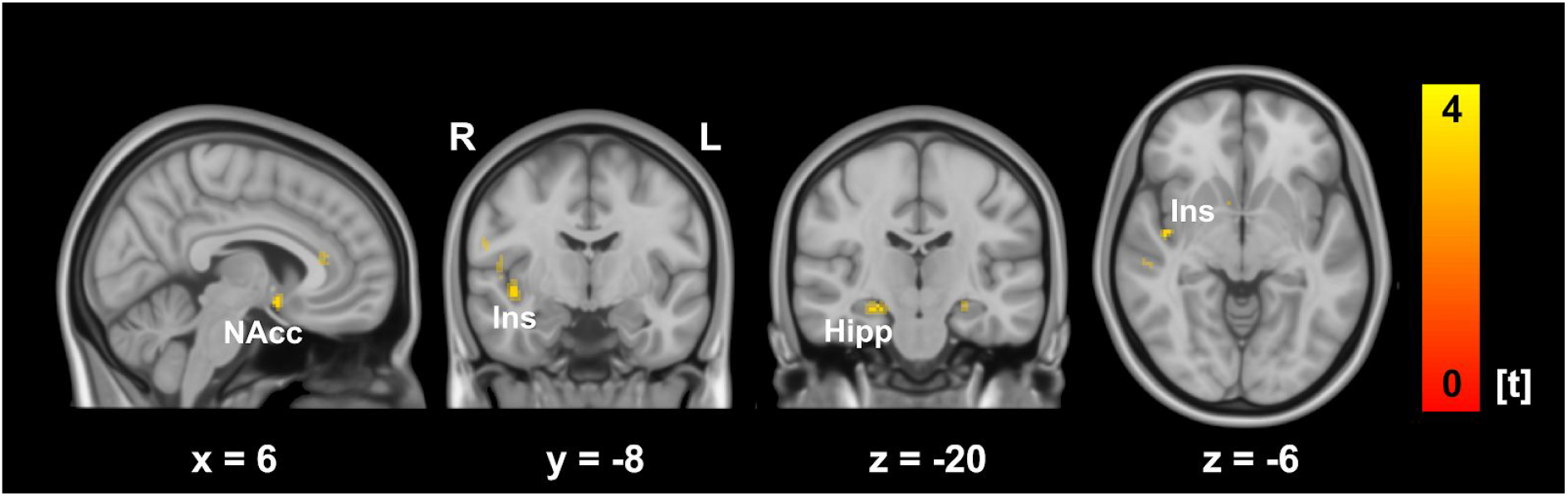
Brain perfusion over the migraine cycle. Brain regions where perfusion is increased with closer temporal proximity to the next attack include limbic areas, particularly the insula, hippocampus and the nucleus accumbens. Bright colours represent a statistical threshold of p<0.001, activities at a lowered threshold of p<0.005 have been included for display reasons in pale colours. NAcc = Nucleus accumbens, Ins = Insula, Hipp = Hippocampus, L = left; R = right.

**Table 2.**
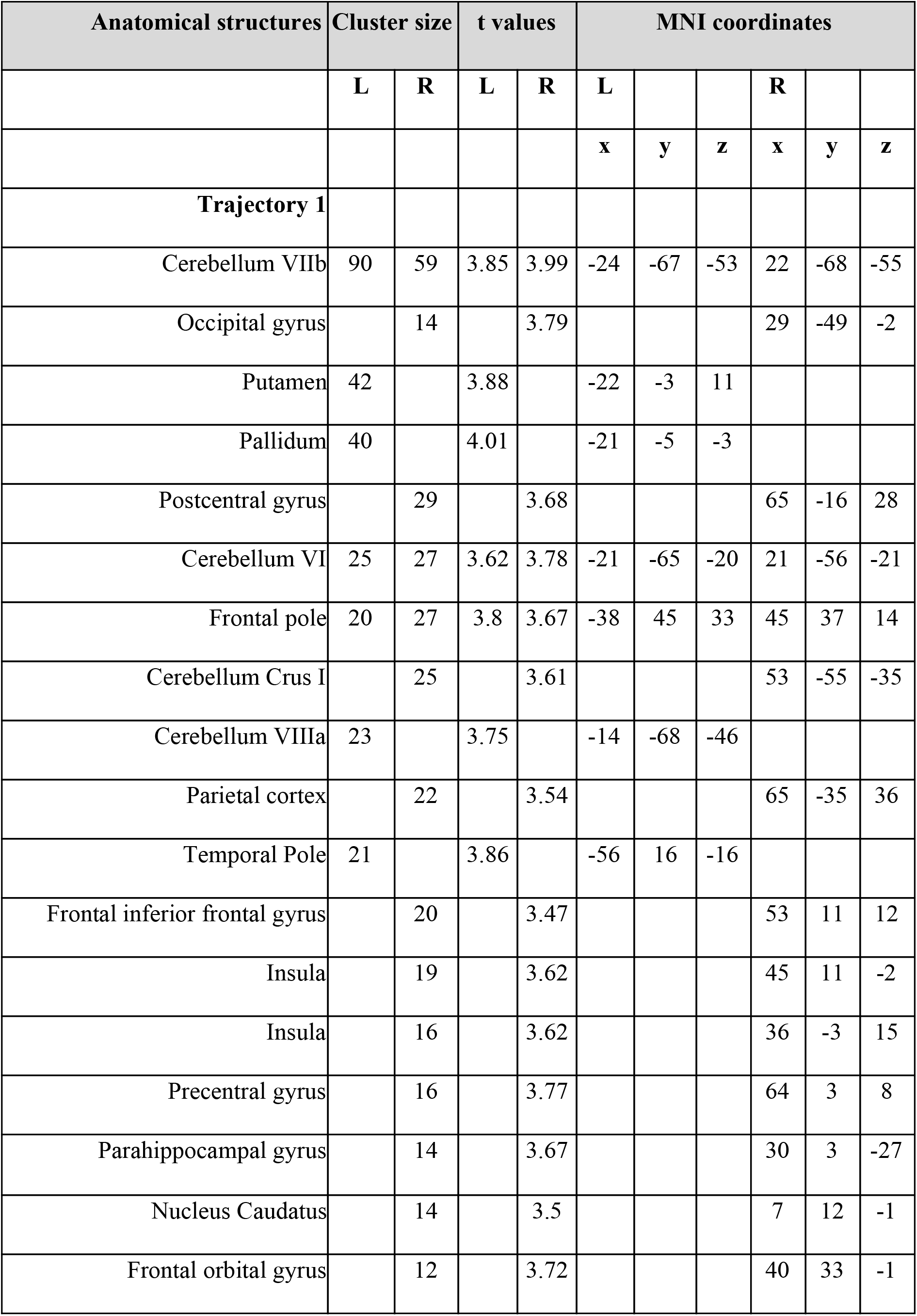
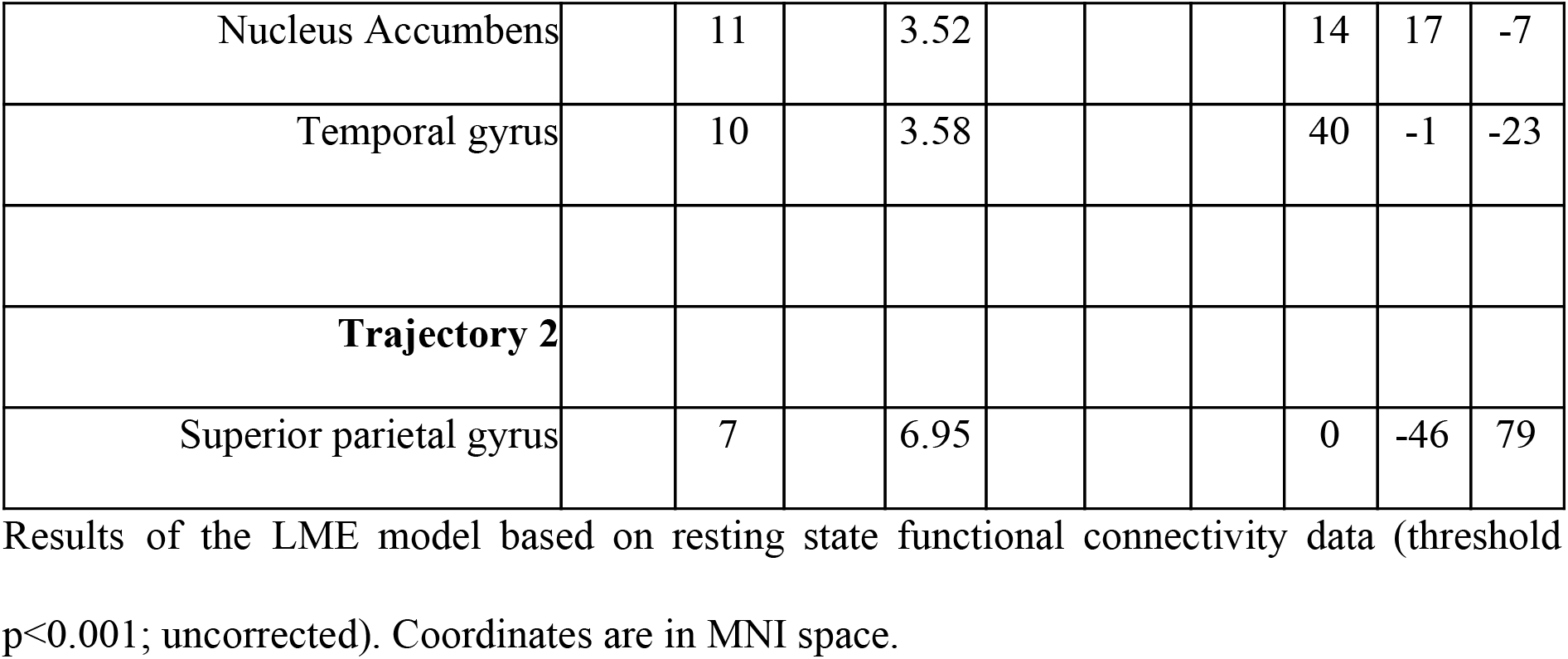
Hypothalamic functional connectivity data over the migraine cycle.

### Hypothalamic functional connectivity

Based on previous research, we tested whether the hypothalamus is changing its functional connectivity over the migraine cycle.

First, we tested for a “drop” of hypothalamic connectivity during headache attacks with a subsequent increase or decrease towards the next attack (“reset mode”). During the attack, we found a low hypothalamic connectivity to the limbic circuit (insula, parahippocampus, nucleus accumbens), the cerebellum, the basal ganglia loop, and sensory, and frontal cortices, which increased towards the next attack (Table 1 and Fig. 2). We did not find any significant effect for the opposite case, for which the hypothalamic connectivity would be high during the attack and steadily decreasing towards the next attack.

**Fig. 2.**
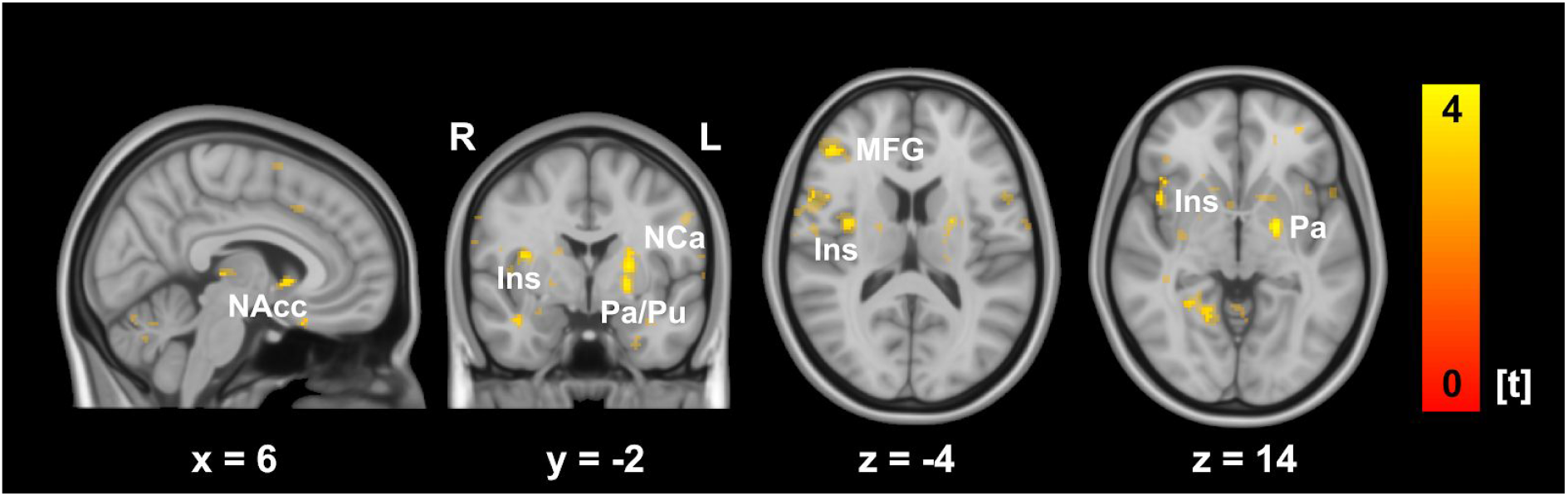
Hypothalamic functional connectivity over the migraine cycle. Brain areas that are functionally connected with the hypothalamus with rhythmic changes over the migraine cycle: These connections are collapsing during the attack and are increasing towards the next attack. This connectivity profile relates to limbic areas, including the insula and the nucleus accumbens, as well as to basal ganglia and frontal areas. Bright colours represent a statistical threshold of p<0.001, activities at a lowered threshold of p<0.005 have been included for display reasons in pale colours. NAcc = Nucleus accumbens, Ins = Insula, Hip = Hippocampus, MFG = Middle Frontal Gyrus, Pa = Pallidum, Pu = Putamen, L = left, R = right.

Next, we tested for hypothalamic connectivity that is changing towards the attack and would reach its climax during the attack (“pain mode”). We found the superior parietal gyrus to follow this trajectory, with the highest connectivity during the attack (Table 2). We did not find any negative trajectory of hypothalamic connectivity throughout the migraine cycle.

## Discussion

The present longitudinal study aimed to detect and record the endogenous and natural rhythmicity of brain activity over an entire migraine cycle by examining episodic migraine patients. Two aspects were specifically addressed: (1) In order to assess the *cyclic behaviour of the brain activity* in migraine patients, we utilised the innovative MRI-based pCASL technique. This allows for voxel-wise measurement of blood perfusion as a proxy for neuronal activity, yet avoids the shortcomings of fMRI BOLD designs requiring a baseline for comparisons. We were able to follow the naturally evolving trajectory of migraine-related rhythmic brain activity without the need of external painful stimulation. We found increasing blood flow over the pain-free interval that reached its maximum during the spontaneous headache attack in the posterior insula, the hippocampus, and the nucleus accumbens. (2) We further mapped the *fluctuating hypothalamic connectivity* across the whole brain over the entire migraine cycle. Here, we observed the connectivity between the hypothalamus and the insula, the nucleus accumbens, and the hippocampus increased over the interictal interval, reaching its maximum shortly before the headache is initiated. In contrast, during the headache this connectivity was low, suggesting a hypothalamo-limbic connectivity collapse. Similar results were found for hypothalamic connections to the basal ganglia (nucleus caudatus, pallidum, and putamen), and cerebellar regions.

### Trajectory of the cerebral blood flow over the migraine cycle

Our longitudinal data over the migraine cycle shows that the perfusion changes occur mainly in limbic structures. We found increasing limbic activity towards the next attack that reached its peak during the migraine attack. This finding most likely reflects a steadily increasing sensitivity of the migraineurs’ brain for sensory stimulation (*23*). The limbic system is a collection of distinct but highly interconnected brain regions that contribute to the processing of fear (*24*), stress reactivity (*25*), learning, and memory (*26*). Via its strong connections to subcortical and prefrontal areas, the limbic system is considered to be a mediator between autonomic reactions and the cognitive evaluation of incoming aversive sensory stimuli (*27, 28*).

In migraine, clinical observations (*29*) and experimental research (*7*) point to a deviant processing of sensory stimuli, which can be attributed to the impaired limbic system. Particularly during attacks, patients report an enhanced sensitivity to sensory input irrespective of the modality (light, sound, odour, and sensation), causing many patients to prefer a dark and quiet environment (*30*). Altered sensory perception, including pain, is not limited to the ictal phase, but has also been observed during the pain-free interval (*23*). Several electrophysiological studies demonstrated a lack of habituation in migraineurs interictally in response to repetitive sensory stimulation (*31*). In contrast, the habituation deficit normalised just before and during attacks (*32, 33*). Due to its important role in integrating incoming stimuli, an impaired limbic system is most likely responsible for the altered sensory processing in migraine patients (*34, 35*). Functional MRI studies revealed enhanced limbic activity during migraine attacks in response to painful (*36*) and olfactory stimulation (*15*). In contrast, hypometabolism of limbic areas, particularly the insula, cingulate and prefrontal cortices, was observed during the pain-free interval, compared to healthy controls (*37*). Other studies observed an altered functional connectivity between the hippocampus, insula, amygdala, and pain-modulating and encoding areas during (*21, 38*) and outside of migraine attacks (*39*).

Further clinical evidence for a migraine-related cyclic trajectory of the limbic system comes from studies on psycho-physiological stress. Processed in limbic structures, psycho-physiological stress is one of the predominant factors for the initiation of the attack (*13*). Stress-related symptoms accompanying the headache, such as reactions of the autonomic nervous system as well as cognitive and attentional deficits, have been linked to limbic processes (*40–42*). Limbic alterations are further related to migraine-related disabilities (*39*), increased pain during attacks (*38*) and the severity of the disease (*37, 43*). These findings offer the opportunity to tailor migraine treatments specifically to limbic functions. Cognitive strategies (*44, 45*), relaxation methods (*46*), and biofeedback (*47*) were shown to be effective techniques to reduce the acute pain sensation and the headache frequency in migraineurs.

Taken together, our findings suggest a cyclic dysfunction of the limbic system. In migraineurs, the limbic system may be genetically determined to be highly vulnerable for sensory input due to ion channel alterations (*48*). The increasing cerebral blood flow in the limbic system over the migraine cycle may reflect an increasing cortical responsiveness to external stimuli. As a result, an impaired limbic function might lead to a general sensory overload in the cortex during the attack, causing a variety of clinical symptoms. The hypothalamus, as a zeitgeber of the limbic circuit, is a candidate for triggering the observed rhythmic changes of the limbic activity.

### Hypothalamic functional connectivity over the migraine cycle

There is strong evidence that the hypothalamus is substantially involved in the pathogenesis of migraine attacks (*7, 8*). Clinical features of the disease correspond to the rhythmic functions of the hypothalamus (*7, 13*). Therefore, we specifically addressed the fluctuations of hypothalamic connections within the brain over the entire migraine cycle. We observed the functional connectivity between the hypothalamus and other limbic centres increased linearly over the interictal interval, reaching its peak shortly before a headache is initiated. In contrast, we found this connectivity to be disturbed during the attack. Similar results were observed for hypothalamic connections to the basal ganglia and cerebellar regions.

The hypothalamus is highly connected with multiple brain regions, including the limbic pathway (*10*). It controls the autonomous and the endocrine system and regulates the haemostatic balance (*11, 12*). These functions are closely related to symptoms during attacks (nausea, vomiting, fatigue, polyuria) and to trigger factors reported by migraine patients, such as changes in hormone status (including menstruation), an unbalanced sleep-wake rhythm, or skipping a meal (*13*). Alterations of the hypothalamus have been observed in previous neuroimaging research: increased activity was detected ictally (*14, 20*) and during the prodromal phase shortly before the onset of pharmacologically-triggered (*19*) and spontaneous headache attacks occurred (*17, 20*). In addition, enhanced hypothalamic-brainstem functional connectivity was observed during migraine attacks, which was already present shortly before the onset of pain. The current finding underpins the importance of the hypothalamo-limbic loop for the generation of migraine attacks. Increasing connectivity between the hypothalamus and limbic areas during the pain-free interval is suggested to reflect the increasing hypothalamic effort to orchestrate limbic subregions that process external and internal information. An insufficient inhibition of limbic excitability in migraineurs may therefore result in a gradual sensory overload of the brain, which is “reset” in a migraine attack. The gradually disrupted hypothalamo-limbic connection suggests that the headache attack is the result of a loss of control by the hypothalamus to limbic structures.

The presented longitudinal MRI study highlights the role of the hypothalamus in generating migraine attacks. Our findings suggest an increasing loss of hypothalamic control over limbic structures, which could result in an increased susceptibility of limbic neurons to the known migraine triggers. Repetitive psychosocial stress, one of the leading trigger factors reported by patients, might make the limbic system even more vulnerable and cause premature triggering of a migraine attack.

Most likely, migraineurs are suffering from a genetically-determined cortical hyperexcitability. As a result, limbic regions in migraineurs may be more sensitive to sensory stimuli. Effects of some prophylactic migraine drugs, including antidepressants and antiepileptics, on the attack frequency and severity are probably relying on the inhibition of these hypersensitive limbic neurons. The hypothalamus as a zeitgeber of internal processes is considered to control the limbic circuits via its strong functional connections. Potential therapeutic interventions are therefore suggested to strengthen limbic circuits with psychological approaches, such as relaxation techniques, cognitive interventions, or biofeedback.

## Materials and Methods

### Subjects

Twenty-two episodic migraine patients were included in the study. Drop-outs from the initially-recruited subjects resulted from incidental MRI findings (n=2), technical problems with the scanner hardware (n=3) or software (n=1), illness of patients (n=2), or taking analgesic medication due to an acute painful event (n=1) during the scanning period. One patient decided to withdraw prematurely from the experiment. In order to collect a sufficient amount of individual data for our longitudinal statistical analyses, we included patients with attack frequencies between 1 and 6 per month. One patient reported a frequency between 6 and 10 attacks per month, this patient was also included but tested daily. Fig. 3 shows the time course of the fMRI recordings for the 12 subjects (11 females and one male). The time points of the attacks were equally distributed in reference to the menstrual cycles of the female patients. Characteristics and clinical features are presented in Table 3. Migraine diagnosis was based upon the classification of the International Headache Society (*30*). The patients did not report any other neurological or psychiatric disorders, were not taking preventative medication for migraine for at least six months, but were allowed to take their regular acute migraine medication after the recording of the headache attack (non-steroidal anti-inflammatory drugs or triptans). Due to their effects on cortical perfusion, patients were not permitted to consume coffee or other caffeinated beverages. All patients gave their written, informed consent. The study was conducted according to the Declaration of Helsinki and approved by the Ethics Committee of the Technische Universität München, Germany. All patients were remunerated for participation.

**Fig. 3.**
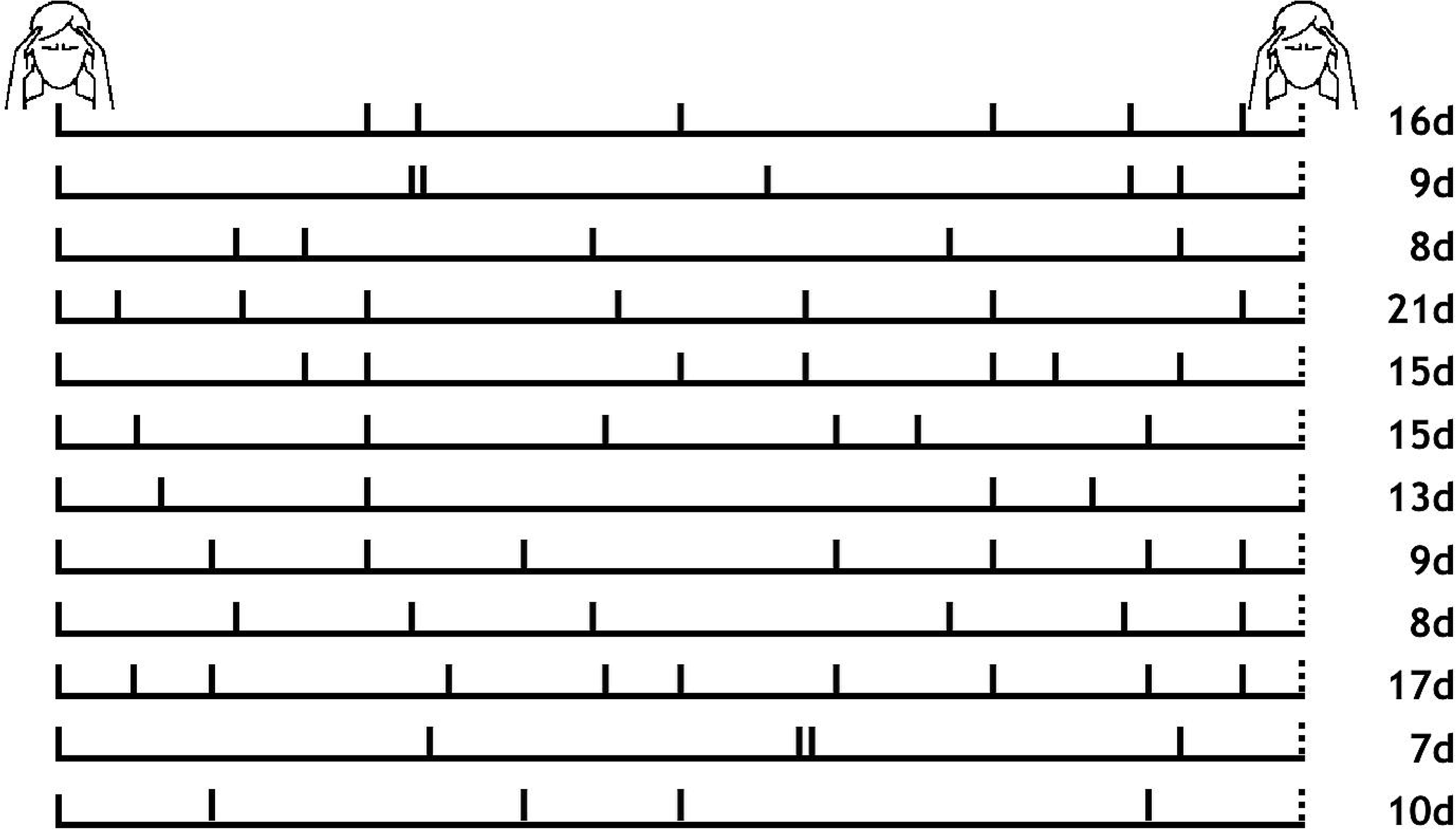
Time course of individual recordings. Solid vertical lines indicate recording days. The dashed line represents the day of the subsequent migraine attack (not recorded) at which the recordings for the subjects were discontinued. The number of days between the first recorded migraine attack and the subsequent migraine attack is shown on the right and indicates the different length of each patient’s migraine cycle.

**Table 3.**
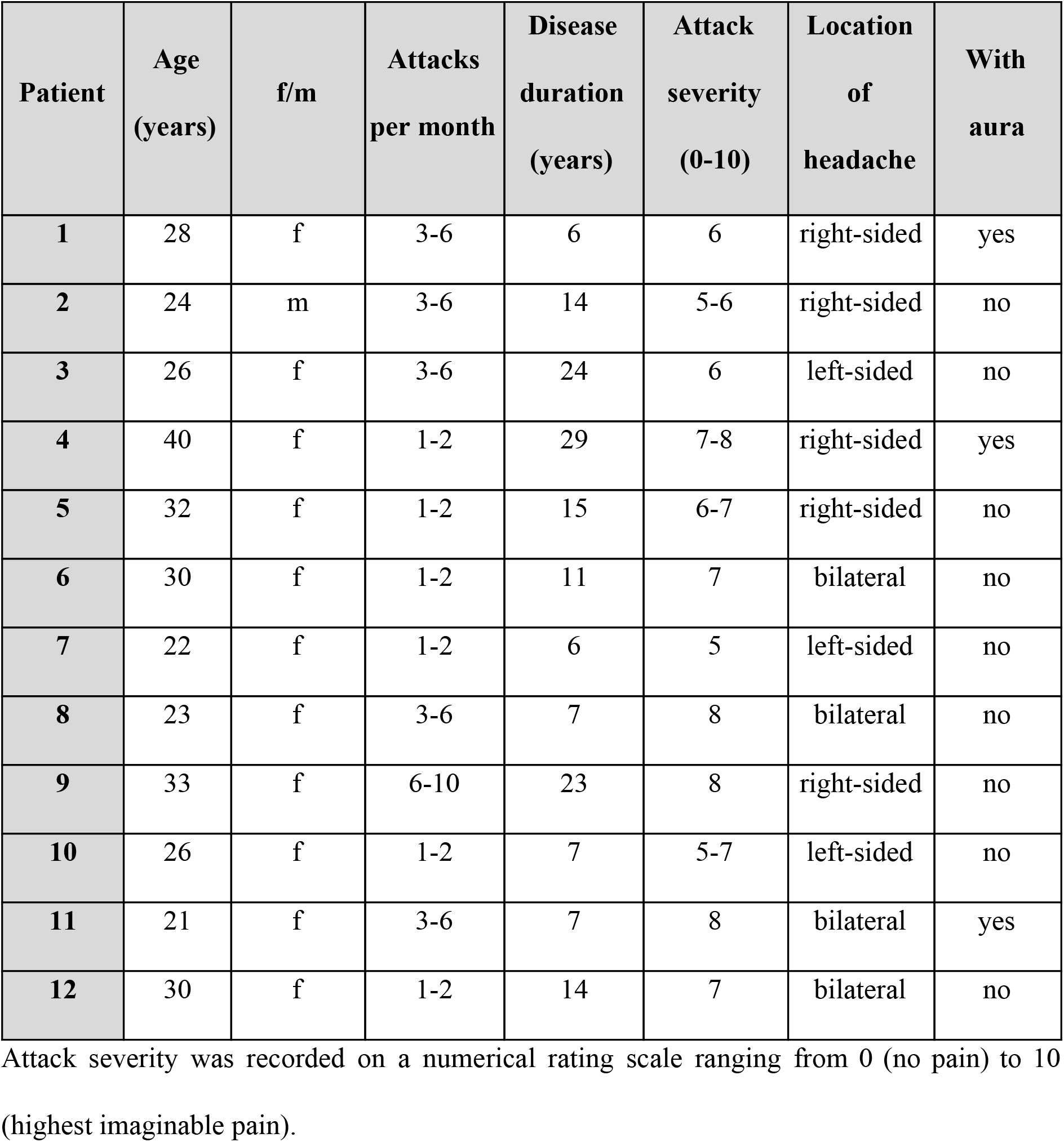
Demographic characteristics and clinical migraine features.

| **Patient** | **Age**<br>**(years)** | **f/m** | **Attacks**<br>**per month** | **Disease**<br>**duration**<br>**(years)** | **Attack**<br>**severity**<br>**(0-10)** | **Location**<br>**of**<br>**headache** | **With**<br>**aura** |
| --- | --- | --- | --- | --- | --- | --- | --- |
| **1** | 28 | f | 3-6 | 6 | 6 | right-sided | yes |
| **2** | 24 | m | 3-6 | 14 | 5-6 | right-sided | no |
| **3** | 26 | f | 3-6 | 24 | 6 | left-sided | no |
| **4** | 40 | f | 1-2 | 29 | 7-8 | right-sided | yes |
| **5** | 32 | f | 1-2 | 15 | 6-7 | right-sided | no |
| **6** | 30 | f | 1-2 | 11 | 7 | bilateral | no |
| **7** | 22 | f | 1-2 | 6 | 5 | left-sided | no |
| **8** | 23 | f | 3-6 | 7 | 8 | bilateral | no |
| **9** | 33 | f | 6-10 | 23 | 8 | right-sided | no |
| **10** | 26 | f | 1-2 | 7 | 5-7 | left-sided | no |
| **11** | 21 | f | 3-6 | 7 | 8 | bilateral | yes |
| **12** | 30 | f | 1-2 | 14 | 7 | bilateral | no |

Attack severity was recorded on a numerical rating scale ranging from 0 (no pain) to 10 (highest imaginable pain).

### Image acquisition

MRI data were collected on a 3 Tesla scanner (Ingenia, Philips, The Netherlands) using a 32-channel head coil. Patients were instructed to remain awake and relaxed with their eyes closed. The imaging parameters of the pseudocontinuous arterial spin labelling (pCASL) data were set according to the latest recommendations (*49*). The labelling plane position was individually planned based on a phase contrast angiography (PCA) survey (scantime 0:21 min). The imaging parameters of the pCASL sequence were: labelling duration = 1650 ms; post label delay = 1700 ms; single-shot 2D EPI readout, repetition time (TR) = 5000 ms, time to echo (TE) = 16 ms; flip angle = 90°, 20 axial slices; field of view (FOV) = 240 mm x 240 mm x 119 mm^3^, voxel size = 2.73 × 2.73 x 5 mm^3^, gap 1 mm, 2 background suppression pulses, 50 repetitions. The total pCASL acquisition time corresponded to 8:30 min. Additionally, a proton density weighted (PDw) M_0_ scan was acquired with TR = 5000 ms and 5 repetitions.

For the 300 volumes of resting state data, we used the following parameters: TR = 2000 ms; time to echo (TE) = 30 ms; FOV = 192 × 192 mm^2^; flip angle = 90°; number of slices = 37; voxel size = 3 × 3 x 3 mm^3^ (0.29 mm gap). For image registration, a high resolution T1-weighted anatomical image was collected with: TR = 9000 ms, TE = 4 ms, flip angle = 8°, FOV = 240 x 240 x 170 mm^3^; number of slices = 170; voxel size = 1.0 x 1.0 × 1.0 mm^3^). Field maps were acquired in each session to control for B0-effects; 64 slices, TR = 960 ms, FOV = 192 × 192 mm^2^; voxel size = 2.0 x 2.0 × 2.0 mm^3^, 0.2 mm gap between slices. TE = 6 ms / 10.55 ms, flip angle 60°.

### Study design

Migraine patients were tested repeatedly over an entire migraine cycle. The imaging time series for each patient started with the recording of a spontaneous and untreated headache attack within the first 6 hours after the beginning of the headache. We only recorded headache attacks which were reported with an intensity of middle to strong, predefined as a minimum of “4” on a numerical rating scale with the endpoints “0” (no pain) and “10” (extremely intense pain). Brain perfusion images were then recorded every 1-4 days at the same time of the day until patients informed us by phone about the following headache attack (which was not scanned) and the time series was completed with the last attack-free recording.

### Data analysis

#### Cerebral blood perfusion (pCASL)

Quantitative cerebral blood flow (CBF) was derived from pCASL. Using SPM12 (*50*), all label and control images were motion corrected, averaged and subtracted to calculate perfusion weighted images. The M0 image was derived from averaging the motion corrected PDw images. By the perfusion weighted images and the M0 image, quantitative CBF-maps were calculated according to recent recommendations (*49*). Those CBF-maps were registered to MNI-space (2 mm isotropic resolution). As we were specifically interested in a very small subcortical region (the hypothalamus), we used a relatively small 3D Gaussian kernel with FWHM of 5 mm for spatial smoothing. Due to occasional artefacts in the vertex region, the analysis of CBF-maps was restricted to axial slices with z-coordinates < 58 mm.

#### Hypothalamic functional connectivity

The data were preprocessed with FSL. The Melodic toolbox was used to execute brain extraction, high-pass filtering with a frequency cutoff of 1/100 Hz, spatial registration to the MNI template, corrections for head motion during scanning, and a spatial smoothing (5mm FWHM). A distortion correction of the images was used based on field maps. The data were further semi-automatically cleaned of artefacts with ICA through Melodic (*51*). The number of components had been automatically estimated by the software and artefact-related components were removed from the data according to the recommendations of Griffanti and colleagues (*52*). Head movement during scanning did not exceed 2mm or 2° in any direction. The hypothalamic spherical region of interest (ROI) was generated according to the MRI atlas of the human hypothalamus (*53*). The ROI was created in MNI standard space and transformed into the native space for each recording session and the mean time course of the hypothalamic activity was extracted. By using the FEAT toolbox, the blood oxygenation level dependent (BOLD) functional connectivity was computed in native space for each voxel by relating the time course of hypothalamus BOLD signal intensity to the time course of all voxels. Connectivity maps were subsequently transformed into MNI standard space for statistical analysis in Matlab.

### Statistical analyses

The statistical analyses are identical for perfusion data and connectivity data. We explored the cyclic change of cortical perfusion and cortical connectivity over the migraine interval. In order to investigate how the cortical processes evolve over the trajectory of measurements, we computed voxel-wise linear mixed effects models (LME) and related the time points within the migraine cycle to the cortical processes of interest. We created a time vector for each patient’s migraine cycle and encoded the day of the recording by assigning numbers between 1 and 2. The numbers 1 or 2 were assigned to the measurement during the headache attack, depending on the following two different trajectories of migraine-related processes in the brain:

**Trajectory 1 (“reset mode”)**: In this model, we aimed to detect the migraine-related activity or connectivity that has its peak (positive or negative) just before the headache attack starts and then *drops to the baseline level (positive or negative) during the headache*. From here, the activity or connectivity linearly increases (or decreases) over the migraine cycle to the next attack. In this hypothetical time course, the brain activity or connectivity on the first day after the attack would be similar to the brain activity during the attack. This trajectory can be interpreted as a cortical “reset” mechanism and is in line with neurophysiological studies reporting a habituation deficit in pain-free migraineurs that normalise during headache attacks (*31*). For 5 measurements over 10 days, the following vector is used to reflect the trajectory 1: ***1*** (=*attack*), ***1.2***, *1.4, 1.6, 1.8*.
**Trajectory 2 (“pain mode”)**: In this model, those migraine-related cortical processes are aimed to be detected that have a *peak during* the headache attack and drop to the level (positive or negative) on the next day. From there, we assume a linear increase (or decrease) over the migraine cycle towards the next attack. We hypothesise increased brain activity in regions that contribute to the processing of migraine symptoms, e.g. pain, increased sensitivity to light, sound, and odors and vegetative complaints. In this hypothetical time course, the brain activity on the day prior to the attack would be similar to the brain activity during the headache attack and is suggested to reflect the increasing excitability of the brain. Similar to the above mentioned example with 5 measurements over 10 days, the following vector is used for the trajectory 2: ***2** (=attack), 1.2, 1.4, 1.6, **1.8***.

The statistical analysis for perfusion and for hypothalamic connectivity has been performed in Matlab (Version R2018a, Mathworks, USA). To explore the relationship between the fluctuating cortical connectivity and the variable pain experience, we computed LMEs (*54*) that related the longitudinal recordings of perfusion and connectivity to the number vector of recording days:

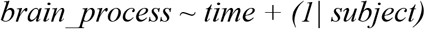

The model is expressed in Wilkinson notation (*55*); the included fixed effect (brain_process ~ time) essentially describes the magnitudes of the population common intercept and the population common slope for the dependency of cortical data on the factor time. The added random effect (e.g. 1 | subject) is used to model the specific intercept differences for each subject. In other words, the model estimates the relationship between the cortical processes in dependence to their occurrence in the migraine cycle (fixed effect). Please note that the statistical terms of fixed effect and random effect are differently used in the common neuroimaging software package SPM as compared with the standard nomenclature in statistics that we are following here. t-values are computed voxel-wise as quotients between the beta estimates and the standard errors of the equation. For a comprehensive introduction into LMEs, see (*56*). The statistical threshold was set to p<0.001 (uncorrected). Patients were only included in the analysis if their movements did not exceed 2 mm in translation and 2° in rotation.

## Author contributions

A.S. and E.S. contributed to the conception and design of the study, acquisition and analysis of the data and drafting the manuscript and figures. L.K. contributed to the acquisition and analysis of the data. M.R. contributed to the acquisition and analysis of the data and drafting the manuscript and figures. S.I. contributed to drafting the manuscript. S.K., C.P. and C.Z. contributed to the acquisition and analysis of the data. V.W. contributed to the analysis of the data. T.R.T. contributed to the conception and design of the study and drafting the manuscript.

